# Toxicity of nitrification inhibitors to soil-living animals

**DOI:** 10.1101/2024.05.29.596422

**Authors:** Marianne Bruus, Paul Henning Krogh

## Abstract

The toxicity of four substances appearing in nitrification inhibitors (DMPP, nitrapyrin, triazole and methylpyrazole) was tested towards three selected soil fauna species, i.e. the springtail *Folsomia candida*, the enchytraeid *Enchytraeus albidus* and the earthworm *Eisenia fetida*. Range-finding and definitive tests were performed according to OECD standard protocols. Test results were compared to expected exposure in order to assess the risk of using nitrification inhibitors. Especially nitrapyrin was estimated to pose a potential risk to soil fauna.

## Introduction

The use of nitrification inhibitors (NI) is a well-established agricultural practice aimed at reducing nitrogen loss after fertiliser application, thereby increasing nitrogen availability for the crop. In practice, however, the effect on plant growth in the form of higher yields is generally modest under Danish conditions (e.g. SEGES 2018, 2020). Nevertheless, studies of different NI’s have found that after fertiliser application, emissions of the potent greenhouse gas nitrous oxide (N_2_O) are significantly reduced (e.g. Akiyama et al. 2010, Qiao et al. 2015, Olesen et al 2028). The growing awareness and ambition to reduce the climate impact of agriculture has led to proposals to use NI enrichment of fertilisers as one of five promising climate instruments for greenhouse gas reduction (Dubgaard and Ståhl, 2018). However, before it is politically decided to promote the use of NI in Danish agriculture, it is necessary to document the potential for nitrous oxide reduction under Danish conditions, and to investigate potential risks to soil organisms and key functions, as well as the fate of the substances in the short and long term, including the risk of leaching of active substances and metabolites. Studies of NI have generally focused on NI as a means of improving N availability for plants, while there is a lack of knowledge about potential side-effects on soil-living organisms. In Denmark, regulation of NI’s is covered by the Order on Fertilisers and Soil Improvers etc. (Landbrugsstyrelsen 2022), and NI’s are thus not subject to the comprehensive environmental studies and assessments as well as ecological risk assessment applicable to pesticides, but are covered by the general rules laid down in REACH (EU 2006). Before widespread use of NI as a climate action agent, any side effects and ecotoxicological effects of NI should be documented and evaluated.

NI’s are a group of substances with very different physio-chemical properties (Subbarao et al., 2006), and therefore it is necessary to document both climate effect and potential side effects for individual NI’s. The active substances are typically added as a coating on commercial fertilizers, or as a liquid for mixing in slurry. This leads to different types of leaching and exposure of soil organisms, and therefore both routes of dispersion should be investigated. Current knowledge about NI’s environmental impact is very limited, and recently 1,2,4 triazole, which is a component of Piadin and one of the NI model substances in this study, has been found in groundwater in Denmark (Rosenbom et al. 2016). Therefore, our investigations will have an immediate and very high level of attention in the community and in the authorities. Most of the efficacy studies with NI have focused on the effects of the substances on nitrogen availability and plant growth, as well as inhibition of nitrifying bacteria such as *Nitrosomonas* (e.g., BASF 1999; Toralbo et al. 2017; Wu et al. 2017). The soil food web of animals and microorganisms is responsible for the degradation of organic matter, including the remineralization of nitrogen, phosphorus and other nutrients, and thus for soil fertility. Ecotoxicological effects and potential positive or negative side effects of NI on soil organisms should therefore be investigated before widespread use. There is a lack of available knowledge about ecotoxicological effects and side effects of marketed NI, both internationally and under conditions relevant to Danish agriculture in terms of climate, soil type and agricultural practices. Preliminary studies of effects of NI on soil microorganisms show that ammonium oxidizing bacteria (target organisms) are inhibited, while other microorganisms, including fungi, do not appear to be significantly affected (Kong et al. 2016, Zhang et al. 2017). The effect of NI’s on soil animals has been very poorly studied, and an ecological risk assessment and studies of side effects are therefore required. Kong et al. (2017) performed a mesocosm experiment with addition of 15N-labeled clover with and without DMPP treatment to a system including the earthworm *Lumbricus terrestris*. Earthworm tissue was analysed after four weeks and showed that DMPP-treated clover was eaten at the same amounts as untreated clover. Earthworm survival was close to 100 %.

This paper presents the basic ecotoxicological studies of NI effects on soil-living animals performed in the KLIMINI project, funded by the Danish Agricultural Agency, Climate Research Program 2019 – 2022 grant no. 33010-NIFA-19-726.

## Methods

Enchytraeids (*Enchytraeus albidus* Henle 1837), springtails (*Folsomia candida* Willem 1902) and earthworms (*Eisenia fetida* Savigny 1826) were selected as test organisms representing soil-living animals.

Due to the lack of existing data on ecotoxicological effects of NI’s, a range-finding test was performed prior to the definitive test. Both types of tests were conducted according to OECD guidelines, i.e. for enchytraeids OECD (2004a), for springtails OECD (2009), and for earthworms OECD (2004b).

For all tests, the soil used was collected at the Foulum experimental station (coordinates 56.495569, 9.568098; sandy loam), dried at 80°C for 24 h, sifted through a 2 mm sieve and re-wetted to a water content of 25 %, based on initial assays to find the appropriate soil humidity.

The NI substances tested were selected as likely candidates for use in Denmark. Four active ingredients were included, i.e., DMPP (≥97% 3,4-dimethyl-1H-pyrazole from Sigma-Aldrich, occurs e.g. as 10 % a.i. in the product Vizura), nitrapyrin (≥98 % 2-chloro-6-(trichloromethyl)pyridine from Sigma-Aldrich, 17,79 % in the product Nlock), triazole (98 % 1,2,4-triazole from Sigma-Aldrich, 2.7-2.9 % in the product Piadin), and methylpyrazole (97 % 3-methylpyrazole from Sigma-Aldrich, 1.4-1.5 % a.i. in the product Piadin).

In the range-finding tests, NI’s were tested at 0, 0.1, 1.0, 10, 100 and 1000 mg a.i./kg dry soil, in 2 replicates. Because nitrapyrin is insoluble in water, it was dissolved in acetone, whereas the other three substances were dissolved in tap water. CuCl_2_ was used as a positive control, at 800 mg/kg dry soil for enchytraeids and earthworms, and at 1000 mg/kg for springtails. The selection of CuCl_2_ concentrations was based on existing data (Caetano et al. 2016, Scott-Fordsmand et al. 2000, Scott-Fordsmand et al. 1997, Amorim et al. 2005, Amorim & Scott-Fordsmand 2012, Lock & Janssen 2002, Posthuma et al. 1997, Spurgeon et al. 1994, Neuhauser et al. 1985, Owojori et al. 2009, Criel et al. 2008). Test conditions are summarised in Table 1.

**Table 1.** Test conditions range-finding tests.

| Species | No. per sample | Soil, g dw per sample | ml water per sample | Duration | Conditions | Endpoints |
| --- | --- | --- | --- | --- | --- | --- |
| *F. candida* | 10 females | 24 | 6 | 21 | 20°C, 16:8 light:darkness | No. adults, no. juveniles |
| *E. albidus* | 10 adults | 20 | 5 | 14 | 20°C, 16:8 light:darkness | No. adults, no. juveniles |
| *E. fetida* | 10 adults | 500 | 125 | 14 | 20°C, 16:8 light:darkness | No. adults, no. cocoons |

Based on the results from the range-finding tests, CuCl_2_ and NI concentrations for definitive tests with the three test species were chosen (Table 2). All tests included 4 replicates of NI concentrations, 6 replicates of controls for NI run as single test, 8 common controls when more NI tests were performed in parallel. Test conditions were similar to those of the range-finding tests, except for duration and endpoints (Table 3).

**Table 2.**
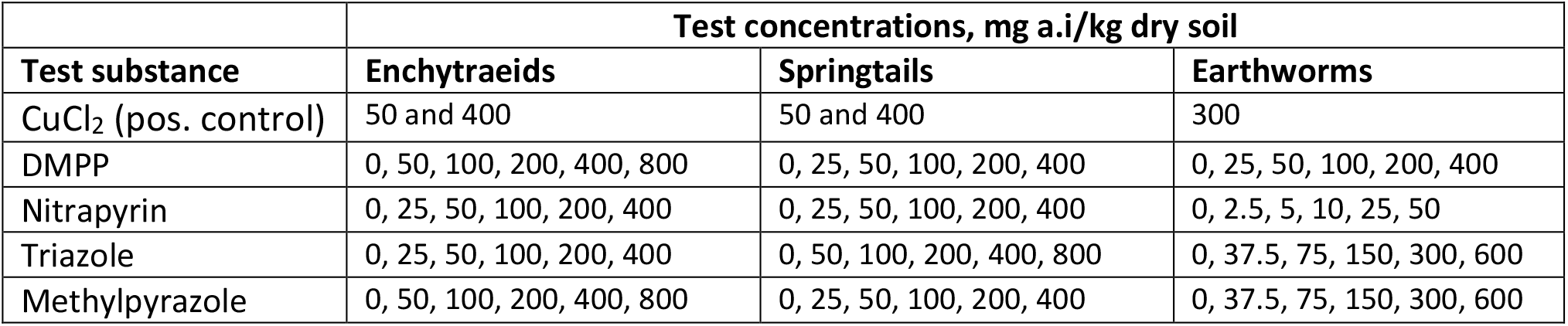
Test concentrations used in definitive tests.

**Table 3.** Test conditions for definitive tests.

| Species | No. per sample | Soil, g dw per sample | ml water per sample | Duration | Conditions | Endpoints |
| --- | --- | --- | --- | --- | --- | --- |
| *F. candida* | 10 females | 24 | 6 | 28 | 20°C, 16:8 light:darkness | No. adults, no. juveniles |
| *E. albidus* | 10 adults | 20 | 5 | 42 | 20°C, 16:8 light:darkness | No adults (day 21), no. juveniles (day 42) |
| *E. fetida* | 10 adults | 500 | 125* | 56 | 20°C, 16:8<br>light:darkness | No. adults (day 28), no.<br>juveniles, no. cocoons (day 56) |

### Statistics

Data from both range-finding and definitive tests were used to estimate effect concentrations by the SAS proc nlin procedure (SAS Institute®). Different dose-response models were used to fit data (cf. Table 4).

**Table 4.** LC_50_ and EC_x_ values [95 % confidence limits] (mg a.i./kg dry soil) estimated on basis of data obtained in range-finding and definitive tests.

| NI | Species | LC<sub>50</sub> range-finding | EC<sub>50</sub> definitive test,<br># juveniles | EC<sub>10</sub> definitive test |
| --- | --- | --- | --- | --- |
| DMPP | *F. candida* | 327 [137-517] | 121 [63-179]<sup>1</sup> | 18 [10-27]<sup>1</sup> |
|  | *E. albidus* | 313 [69-557] | 105 [68-141]<sup>2</sup> | 21 [14-28]<sup>2</sup> |
|  | *E. fetida* | 290 [144-438] | 132 [-12-278]<sup>3</sup> | 20.1 [-1.9 - 42.2]<sup>3</sup> |
| Nitrpyrin | *F. candida* | 207 [55-359] | 300 [263-337]<sup>4</sup> | 220 [212-227]<sup>4</sup> |
|  | *E. albidus* | 148 [60-237] | 16 [6-26]<sup>1</sup> | 2.5 [0.93-4.0]<sup>1</sup> |
|  | *E. fetida* | 108 [-19-235] | 11 [0.35-22]<sup>1</sup> | 1.7 [0.05-3.4]<sup>1</sup> |
| Triazole | *F. candida* | 2500 [984-4015] | 441 [356-525]<sup>5</sup> | 283 [146-420]<sup>5</sup> |
|  | *E. albidus* | 50 [5.2-95] | 45 [10-81]<sup>1</sup> | 6.9 [1.4-12]<sup>1</sup> |
|  | *E. fetida* | No effects within tested range | 85 [59-112]<sup>1</sup> | 13 [9-17]<sup>1</sup> |
| Methyl-<br>pyrazole | *F. candida* | 322 [147-498] | 195 [145-246]<sup>6</sup> | 114 [107-122]<sup>6</sup> |
|  | *E. albidus* | 325 [115-535] | 247 [178-317]<sup>7</sup> | 129 [116-143]<sup>7</sup> |
|  | *E. fetida* | 322 [133-512] | 202 [78-326]<sup>1</sup> | 31 [12-50]<sup>1</sup> |

1 $c \cdot e^{-a \cdot \text{conc}}$ , exponential decrease
2 $c - \text{conc} \cdot a$ , linear decrease
3 $c \cdot e^{-a \cdot \text{conc}}$ , exponential decrease; the definitive test failed, # cocoons from the range-finding test was used for EC<sub>10</sub> and EC<sub>50</sub>.
4 $c + a \cdot (\text{conc} - 200)$ , linear decrease; no effects below 200 mg/kg.
5 $c / (1 + (a \cdot \text{conc})^b)$ , sigmoid
6 $c \cdot e^{-a \cdot \text{conc}}$ , exponential decrease starting at 100 mg/kg
7 $c - \text{conc} \cdot a$ , linear decrease starting at 100 mg/kg

## Results

In the range-finding tests, LC_50_ values were determined for all combinations of test species and test substance, except that triazole had no effects on earthworm survival within the range of concentrations tested (Table 4).

**Table 4.**
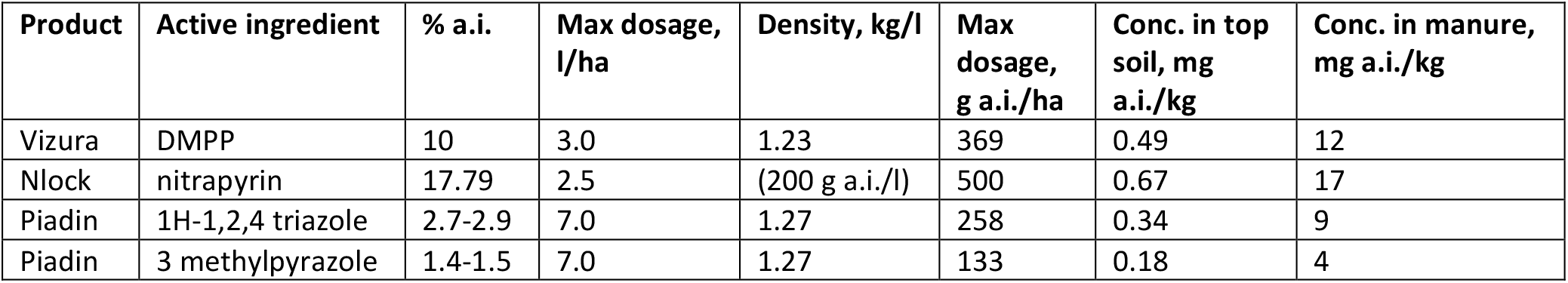
Estimated exposure to NI’s, assuming maximum dosage and distribution in the top 5 cm of the soil. The last column presents the estimated concentrations in manure applied at 30 tons per hectare.

| Product | Active ingredient | % a.i. | Max dosage, l/ha | Density, kg/l | Max dosage, g a.i./ha | Conc. in top soil, mg a.i./kg | Conc. in manure, mg a.i./kg |
| --- | --- | --- | --- | --- | --- | --- | --- |
| Vizura | DMPP | 10 | 3.0 | 1.23 | 369 | 0.49 | 12 |
| Nlock | nitrapyrin | 17.79 | 2.5 | (200 g a.i./l) | 500 | 0.67 | 17 |
| Piadin | 1H-1,2,4 triazole | 2.7-2.9 | 7.0 | 1.27 | 258 | 0.34 | 9 |
| Piadin | 3 methylpyrazole | 1.4-1.5 | 7.0 | 1.27 | 133 | 0.18 | 4 |

The definitive test resulted in estimates of EC_10_ and EC_50_ for reproduction (Table 4), except for effects of DMPP on earthworms, which failed due to soil being too wet; the water content was identical to the other tests, but a new batch of soil was applied this final test, which apparently had a slightly different water holding capacity. As a substitute for the missing data, reproduction data from the range-finding test are given.

### Expected exposure to NI

NI’s may be applied together with mineral fertilisers or manure, for instance pig slurry. In order to assess the risk of NI application for soil living animals, effects levels must be compared to expected exposure levels. If NI is applied with mineral fertilisers, we assume that the NI is mainly distributed in the top 5 cm of the soil (Table 4), whereas application with manure will probably result in a lower soil concentration, unless the manure is incorporated in the top soil, which is rather likely in Danish agriculture. In that case, the worst-case scenario is assessed to be a distribution in the top 5 cm of the soil. Since some soil animals may be exposed in the manure, the maximum concentration in this substrate is also presented.

### Tentative risk assessment

Following the common practice for the risk assessment of pesticides (European Commission 2002), we have established TER’s (Toxicity/Exposure Ratios), i.e. ratios between estimated EC_10_ values and expected exposure, primarily on basis of results from the definitive tests (Table 5). EC_10_ values from the definitive tests were used as a proxy of no effect levels, which are traditionally used to calculate reproduction TER’s. For the effect of DMPP on earthworms, reproduction EC_10_ could not be established from the definitive tests, and therefore the EC_10_ value from the range finding test was used as substitute.

**Table 5.** TER’s for estimated exposure via soil or slurry. established by combining estimates from Tables 3 and 4. Figures in bold indicate TER’s below the criterium (TER<5) that would trigger field testing in the risk assessment of plant protection products according to European Commission (2002).

| Active ingredient | Toxicity/exposure ratios |  |  |  |  |  |
| --- | --- | --- | --- | --- | --- | --- |
|  | Springtail reproduction |  | Enchytraeid reproduction |  | Earthworm reproduction |  |
|  | Soil | Slurry | Soil | Slurry | Soil | Slurry |
| DMPP | 38 | **2** | 43 | **2** | 41* | **2\*** |
| nitrapyrin | 328 | 13 | **4** | **0.1** | **3** | **0.1** |
| 1H-1,2,4 triazole | 832 | 31 | 20 | **1** | 38 | **1** |
| 3 methylpyrazole | 633 | 38 | 719 | 43 | 172 | 10 |

\* Estimates based on EC<sub>10</sub> from range-finding tests.

## Discussion

The observed effects on the reproduction of soil animals after 28-56 days were compared with the expected exposure doses, either via manure (30 t/ha) with maximum doses of nitrification inhibitors or by mixing the nitrification inhibitors into the top 5 cm of the cultivation soil in order to establish TER’s. In risk assessment of pesticides, a TER for effects on reproduction of less than 5 gives rise to further investigations or risk-reducing measures (European Commission 2002) and is therefore an expression of potentially negative effects. If the same threshold value were used for nitrification inhibitors, which is not required, the exposure in manure would exceed the value for DMPP, nitrapyrin and 1H-1,2,4-triazole, while only nitrapyrin appears to be problematic for soil fauna upon exposure in the top 5 cm of the cultivation soil.

The exposure in the laboratory tests cannot be directly compared to the use of nitrification inhibitors in the field, where the substances are applied after mixing with either slurry or commercial fertiliser. Soil animals are expected to avoid direct contact with the anaerobic and ammonia-containing environment in newly added manure (e.g. Curry 1976), whereas after a few weeks manure is attractive to soil animals and stimulates population growth (e.g. Curry 1976, Silva et al. 2016). Therefore, the greatest potential for exposure is expected to be when applied directly to the soil together with commercial fertilizers or via slurry that has been on the soil/in the soil for a few weeks, where the concentration of nitrification inhibitors is expected to be reduced due to partial decomposition (Byrne et al. 2020). However, if nitrification inhibitors are applied several times during the growing season, the risk of negative effects on soil animals may increase, depending on whether the substances break down before the next application.

In line with the present study, previous studies (e.g. Kong et al. 2017, Bachtsevani et al. 2021, Rodrigues et al. 2018 and Schmidt et al. 2022) have shown that the different active substances and formulations of nitrification inhibitors have very different properties and potential effects in nature. Nitrification inhibitors are added together with a nitrogen source, and their distribution and interaction with soil-living organisms is thus linked to the turnover in nutrient-rich environments. This complicates the risk assessment, and there is a need for practical information about the impact of the various nitrification inhibitors on soil-living organisms and water quality.

